# MIADE metadata guidelines: Minimum Information About a Disorder Experiment

**DOI:** 10.1101/2022.07.12.495092

**Authors:** Bálint Mészáros, András Hatos, Nicolas Palopoli, Federica Quaglia, Edoardo Salladini, Kim Van Roey, Haribabu Arthanari, Zsuzsanna Dosztányi, Isabella C. Felli, Patrick D Fischer, Jeffrey C. Hoch, Cy M Jeffries, Sonia Longhi, Emiliano Maiani, Sandra Orchard, Rita Pancsa, Elena Papaleo, Roberta Pierattelli, Damiano Piovesan, Iva Pritisanac, Thibault Viennet, Peter Tompa, Wim Vranken, Silvio CE Tosatto, Norman E Davey

## Abstract

An unambiguous description of an experimental setup and analysis, and the subsequent biological observation is vital for accurate data interpretation and reproducible results. Consequently, experimental analyses should be described in a concise, unequivocal, and digestible manner. The aim of minimum information guidelines is to define the fundamental complement of data that can support an unambiguous conclusion on experimental observations. In this document, we present the Minimum Information About Disorder Experiments (MIADE) guidelines to define the minimal fundamental parameters required for non-experts to understand the key findings of an experiment studying intrinsically disordered proteins (IDPs) or intrinsically disordered protein regions (IDRs). MIADE guidelines provide recommendations for data producers to describe the results of their experiments at source, for curators to annotate experimental data to community resources and for database developers maintaining community resources to disseminate the data. We give examples of the application of these guidelines in common use cases and describe the implementation of an update to the DisProt IDP database to allow MIADE-compliant annotation. The MIADE guidelines will improve the interpretability of experimental results for data consumers, facilitate direct data submission, simplify data curation, improve data exchange among repositories and standardise the dissemination of the key metadata on an IDP experiment by IDP data sources.

## Introduction

The intrinsically disordered protein (IDP) field is generating increasingly large amounts of biophysical data on the structural properties of intrinsically disordered regions (IDRs)^1,2^. The complexity of the produced IDP-related data continues to increase, and in recent years there has been a noticeable growth in the number of analyses describing complex structural properties, conditional disorder and disorder-function relationships^3–6^. Whereas a decade ago most IDP papers characterised disorder as a binary state, now many papers contain comprehensive analyses describing multiple conditional states using several complementary experimental methods^7,8^. Moreover, the improved experimental tools now enable the investigation of increasingly complex IDRs, IDPs, and multi-domain proteins. A key responsibility of the IDP community is the development of minimum information guidelines to improve the description, interpretation, storage and dissemination of data generated in the rapidly evolving IDP field^9^. In this document, we introduce the Minimum Information About Disorder Experiments (MIADE) guidelines for the definition and interpretation of experimental results from IDP experiments.

Minimum information guidelines define the fundamental unit of information for the unambiguous definition of experimental metadata to the level required for a non-expert to comprehend the key results of an experiment^10^. The role of minimum information guidelines is to minimise data loss by preserving essential data and removing ambiguity while avoiding redundancy. There are several requirements for a functional minimum information guideline. Firstly, the core information conveyed by the experiment should be unequivocally defined. This should include the observation itself but also any information that would change our understanding or confidence in the biological or physical relevance of the observation. Second, adhering to the guidelines should be as effortless as possible to enable its widespread adoption, i.e., the guidelines should avoid any excessive burden in the description of an experiment while capturing the most important information to fulfil the first requirement. Thirdly, it should be equally applicable to all IDR analysis methods so that the experimental metadata is comparable across all sources of primary data, regardless of the experimental approach. To fulfil these criteria, the MIADE guidelines recommend an unambiguous description of the protein and the construct of the region(s) being studied at amino acid resolution, other components of the sample, the experimental approach applied and the expert interpretation of the results of the experiment. Importantly, any information about the experimental protocols, sample components or sequence properties that might affect the interpretation of the results are an essential part of the unambiguous description of the experimental results.

Minimum information guidelines are a compromise between the necessary depth of information to unambiguously describe an IDP experiment, and the reporting burden placed on researchers producing the metadata. MIADE-compliant data records should allow users to quickly assess an IDP experiment and the associated data and point to the source data for the complete experimental context, but do not require annotation to a level of detail that allows the experiment to be reproduced. Therefore, unless their definition is essential to unambiguously interpret the results of the experiment, several aspects of the experimental setup are not required by the MIADE guidelines; this includes a complete description of the experimental constructs, a complete description of the sample and a complete description of the experimental setup. Furthermore, minimum information guidelines are abstract recommendations that do not specify the technical details of the structured data types that are guideline compliant. In this document, we provide examples of data adhering to MIADE recommendations for multiple use cases including providing details of the updates to allow the storage of MIADE-compliant data in the DisProt IDP database^1^. However, the technical specification of data storage is defined by exchange formats used to standardise and store compliant data and therefore it is outside the scope of this document.

The MIADE guidelines provide a community consensus created by experimentalists, curators and data scientists on the minimum information required to appropriately describe metadata on experimentally and computational-derived structural state(s) of IDPs or IDRs. The aim is to increase the accuracy, accessibility and usability of published IDP data, to comply with FAIR (findability, accessibility, interoperability, and reusability) data principles^11^, to support rapid and systematic curation of such IDP data in public databases and to improve interchange of IDP data between these IDP resources. We believe that these guidelines will provide an important roadmap to the thousands of data producers, curators and database developers in the IDP field and increase the utility of published IDP data for the larger biological community.

### Where should MIADE be applied?

The vast majority of IDP experiments yield information about the structure or the function of IDPs. Functional IDP studies most commonly analyse their interactions with other molecules. Since the Minimum Information about a Molecular Interaction experiment (MIMIx) guidelines^12^, on which the MIADE guidelines have been modelled, already cover the molecular interaction aspects of these experiments, MIADE only focuses on the description of the structural aspects of the studied IDPs.

Structural IDP data can follow many paths to the final data consumer (Figure 1A). At each point in the flow of data valuable information can be lost, misinterpreted, or misrepresented. After data production, the primary data are analysed by field-specific experts (typically the research group that conducted the experiment) who interpret these complex experimental results to provide a biological observation. These experts will author a publication that describes the novel observations and, ideally, they will directly submit the findings to a core IDP data resource. Currently, much of the data in the IDP field passes into a branch where biocurators interpret the description of the experiments and observations in the publication and then annotate the information into manually curated resources. The role of MIADE is to provide general recommendations that can be applied at each potential point of data loss to maximise the precision with which information is transferred.

**Figure 1.**
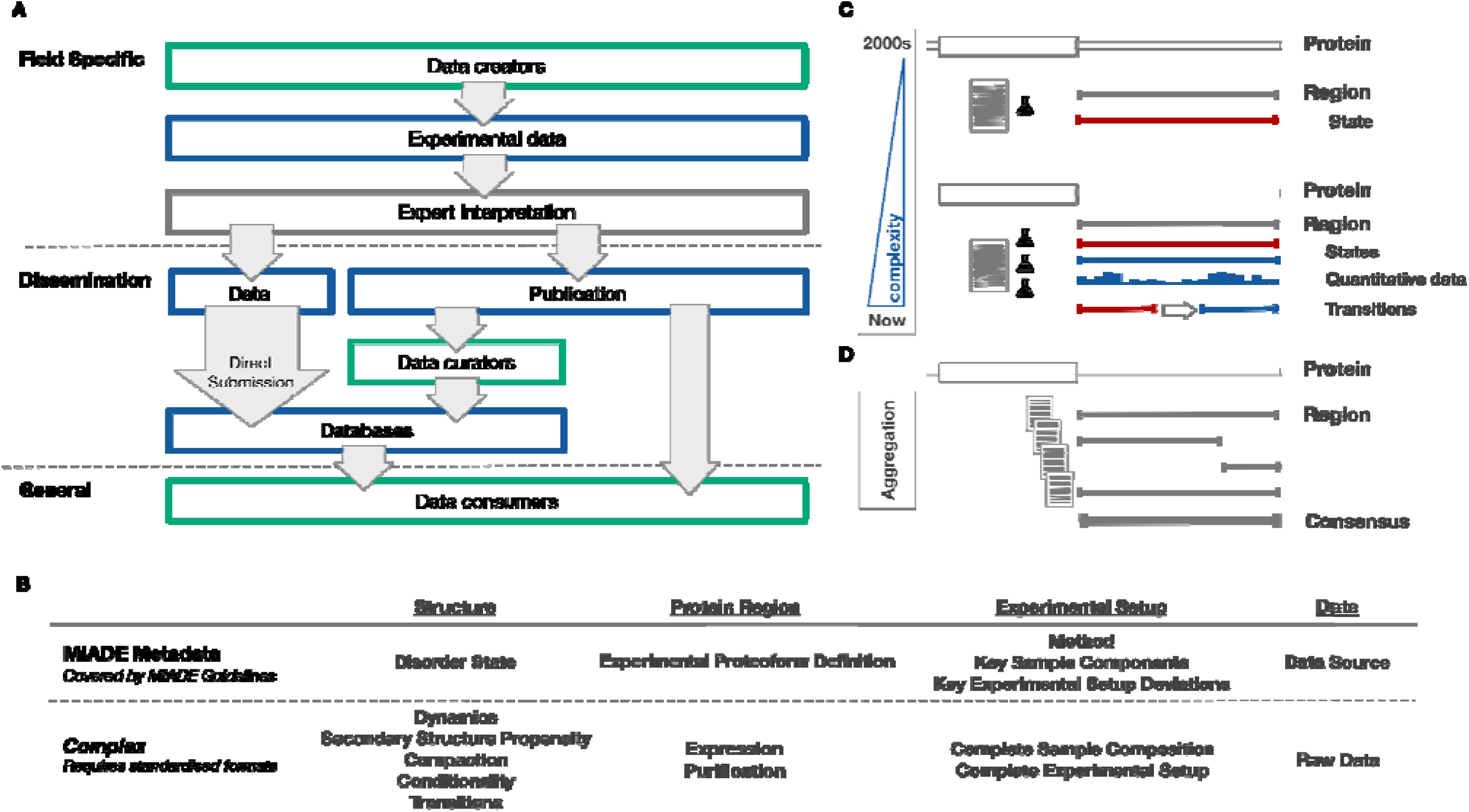
(A) Scheme of data flow from primary data capture by the experimentalist to data dissemination to the end consumer. (B) Definition of the scope of the MIADE guidelines and the requirements of a comprehensive standard for IDP data. (C) Representation of the evolution of complexity of cutting edge experimental IDP papers. (D) Representation of the requirement for data aggregation across analyses to build high confidence consensus data on a region.

The MIADE guidelines should be applied to free text descriptions when reporting on the performed experiment, to data extraction from the primary literature and to structured metadata for dissemination. Therefore, the MIADE guidelines provide a recommendation to unambiguously describe structural information on IDRs inferred from experimental or computational analysis, intended for: (i) researchers authoring an article on the structural state(s) of an IDR; (ii) researchers who want to directly submit such data to an IDP resource, e.g. prior to peer reviewed publication of the data; (iii) biocurators who want to define/curate data on structural state(s) of an IDR within an IDP resource; (iv) database developers who want to disseminate IDR structural state data; and (v) data users who need to achieve full comprehension by clarity of the meaning and origin of each piece of data (Table 1).

**Table 1.** Cases where the MIADE guidelines should be applied to improve data interpretability and minimise the loss of key data.

|  |  |
| --- | --- |
| **Storing experimental metadata** | - Allows storage of high-level metadata.<br>- Allows the integration and comparison of data from distinct experiments and experimental approaches. |
| **Direct submission of IDR data pre-publication** | - Promotes early data capture by providing a standard with a low barrier for data entry to directly submit experimental results prior to publication to an IDP database.<br>- Facilitates collection of IDR data in light of increasing data management and open science efforts.<br>- Increases data available for community blind testing of computational IDP tools (CAID). |
| **Defining key findings about IDRs in a publication** | - Defines the requirements for unambiguous description of the expert interpretation of primary source data.<br>- Increases the clarity of the paper and simplifies all downstream data interpretation.<br>- Allows the reported structural IDR data to be rapidly captured in an IDP resource, where it is readily available to the community. |
| **Curation of IDRs from a publication** | - Provides clear guidelines to unambiguously and efficiently define the fundamental details and results of the experiment(s) to facilitate data curation by non-experts. |
| **Transfer of IDR data** | - Standardised transfer of the key metadata on an IDR experiment to, between, and from IDR data repositories.<br>- Promote/facilitate the implementation of FAIR principles with the IDR community. |

### What information is required by MIADE guidelines

Both the biological and the methodological contexts are required to understand and compare experimental data. Consequently, MIADE guidelines recommend the clear definition of four components for IDP structural experiment reporting: the protein region that was studied, the structural state of that region as inferred from the experiment, the experimental or computational approach applied and the data source. Each region of a protein for which a structural state was inferred from an experiment should be described separately. The exact application of the guidelines is use case specific, however, when possible stable identifiers of external resources should be referenced, for example, UniProt for protein definitions^13^, ECO (Evidence and Conclusion Ontology) for experimental definitions^14^ and IDPO (Intrinsically Disordered Proteins Ontology) for structural state definitions (https://disprot.org/ontology).

#### MIADE Checklist - minimising ambiguity in the definition of a disorder experiment

The following information is required to create MIADE compliant description of an IDP experiment:

##### Protein Region

definition of the region for which a structural state was experimentally determined or computationally predicted. If several regions of a protein were inferred to be disordered, each region should be defined separately. The definition should be unambiguous and concise, and should leave no doubt about the identity of the protein that contains the region. The source organism and isoform should always be specified. If the sequence is synthetic and not mappable to an existing protein this should be stated explicitly. The experimental sequence of the protein region being studied should always be defined. Similarly, any tags, labels, post-translational modifications or mutations present in the sample under study should be described. Each region should be characterised by:

- Definition of the source protein from which the region was derived:

- The common name for the source molecule. Both the protein name and gene name should be added whenever possible. Ideally, this should be the official name provided by a nomenclature committee such as the HGNC symbol from the HUGO Gene Nomenclature Committee for human genes^15^. In cases where the field-specific name is used, and it differs from the official name, the official name should be mentioned in the first definition of the molecule. *Example: Mitotic checkpoint serine/threonine-protein kinase BUB1 beta (BUBR1, also known as BUB1B)*
- Scientific name, common name or NCBI taxonomy ID of the species of origin for the source protein (or free text for chemical synthesis, unknown, and *in silico* origins). *Example: Budding Yeast (Saccharomyces cerevisiae strain ATCC 204508 / S288c, NCBI Taxon ID: 559292)*
- Accession or identifier for the source protein in a reference database. If an isoform of a protein was used in the experiment, the accession or identifier specifically identifying that isoform should be used whenever possible. The version number of the protein sequence in the database can be added to further reduce ambiguity. *Example: UniProt:P13569 (P13569-2 in case isoform 2 was used)*

- Definition of the protein region(s) for which a structural state was determined:
- Start and stop positions of the region: the position of the first and last residue of the region, based on i) the sequence as described in the database annotating the source protein from which the region was derived (i.e. positions should refer to the natural sequence and should not consider added purification and solubility tags), or on ii) in the case of a sequence that is not mappable to a natural sequence, the sequence provided by the data producer. *Example: residues 708-831 of BUBR1*

- The amino acid sequence of the experimental construct encoding the region(s) in IUPAC one-letter codes^16^.
- Definition of the experimental molecule (i.e. any tags in the construct that have been removed before the sample has been studied can be ignored) including any alterations and additions to the defined protein region
- Tags and labels that are present in the experimental construct. *Example: C-terminal 6xHis tag*
- Experimental proteoform including mutations or modifications. *Example: phosphorylation of BUBR1 on serine 21*

##### Structural state

structural state of the construct or a region(s) within the construct, as defined by the experimental data or as inferred by the experimentalist.

- Classically, structural states would be defined as “order” and “disorder”, however, more complex structural properties are now being experimentally defined. The position of a structurally distinct subregion of a construct, such as the observation of partially populated secondary structural elements, should be defined explicitly as described for the protein region definition. If the boundaries of the structure state elements within a construct are not clear this should be stated. When possible, the corresponding term and term ID for that structural state in the IDPO controlled vocabulary should be given. If a structural property that is not widely known to non-experts is used, the term should be clearly defined. *Example: disorder (IDPO:00076)*

##### Experimental and computational approaches

definition of the experiment or computational approach used to determine the structural state of the region. Each experimental setup should be described separately, using the following parameters:

- The experimental or computational methods used to determine the structural state of the region. If possible, this should be annotated with the corresponding term and term ID for that experimental method in the ECO controlled vocabulary. The name of the computational or experimental method(s) used to define the structural state of the protein region(s) should be defined to the most detailed level possible. If relevant, any software used in the post-processing of experimental data, or to define the structural state directly, should be defined including the software version. *Example: far-UV circular dichroism (ECO:0006179)*
- The scientific name, common name or NCBI taxonomy ID of the host organism in which the experiment was performed (or free text for *in vitro*, unknown, *in vivo*, and *in silico* experimental environments); further specification of cell line or tissue is recommended. Special care should be taken in defining experimental details for in-cell or cell extract studies. *Example: in vitro*
- Any experimental deviation that could alter the interpretation of the results and any condition that could impact on the results should be clearly described. These deviations are generally method specific, for example, *in vitro* experimental parameters (i.e. pH, pressure, protein concentrations, temperature, buffer, salt, additional components including other proteins), computational parameters (non-default options) and Molecular Dynamics (MD) simulation parameters (force field used). See next section and Table 2 for details. *Example: experiment was performed at 4°C*.

**Table 2.**
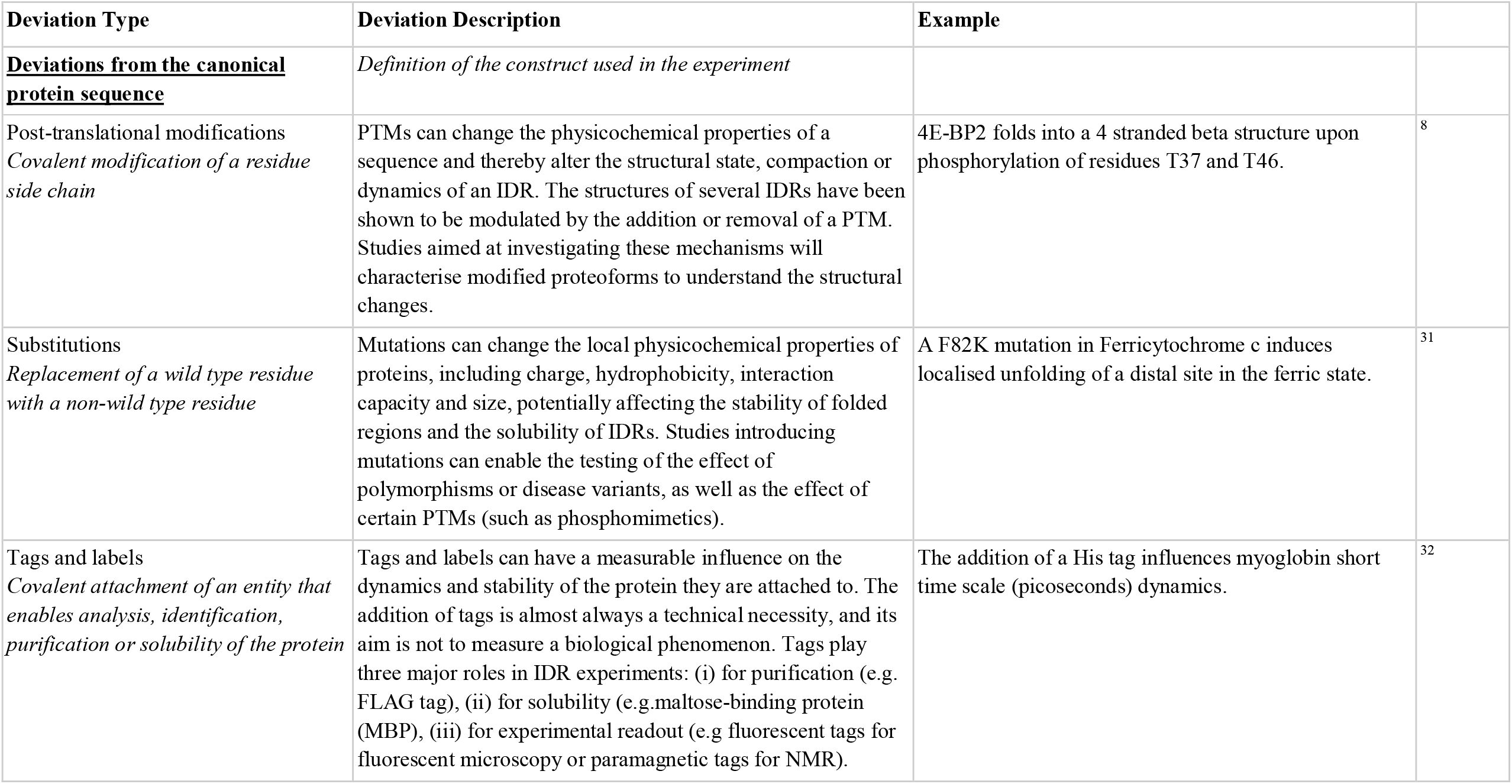
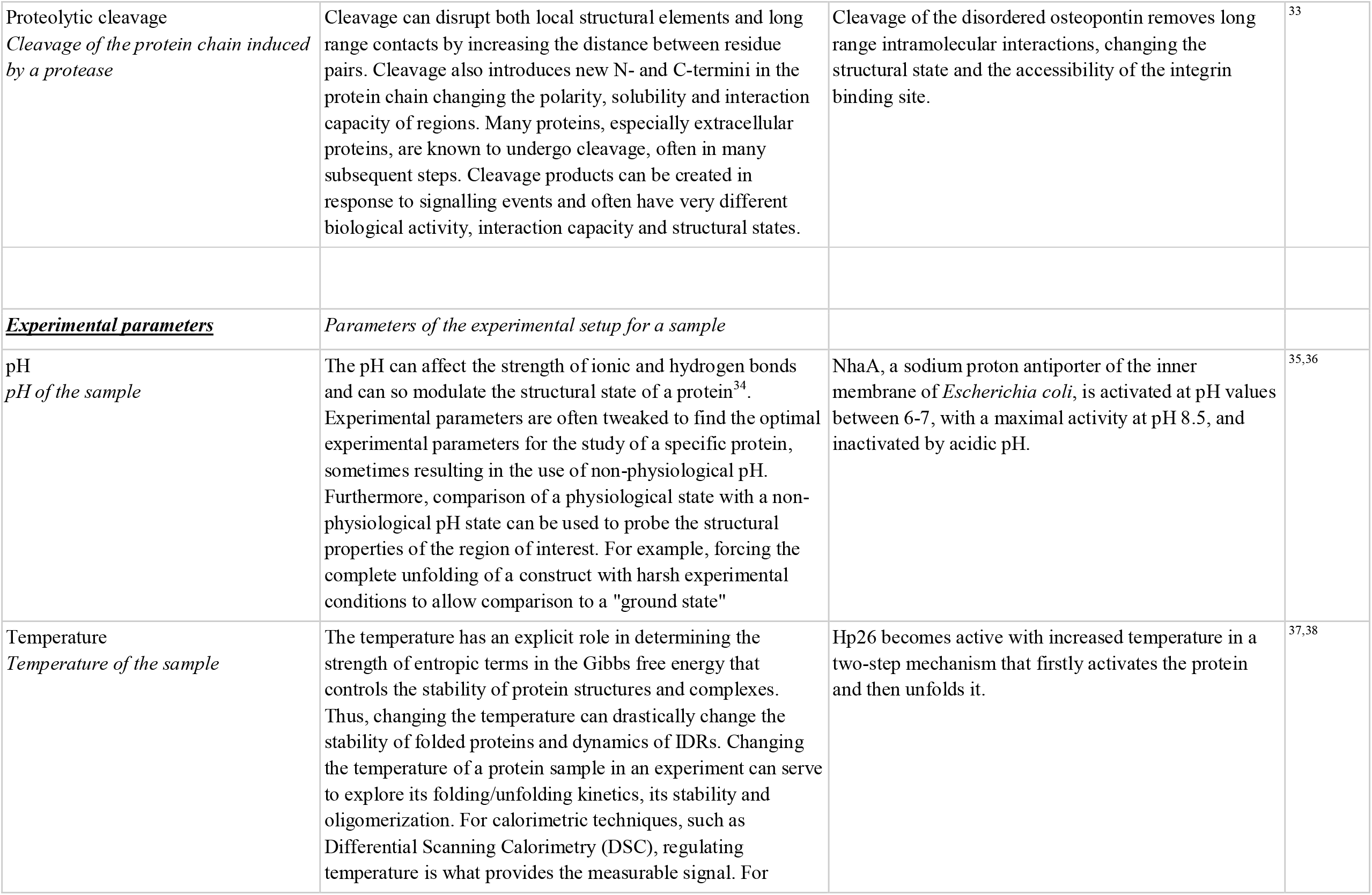
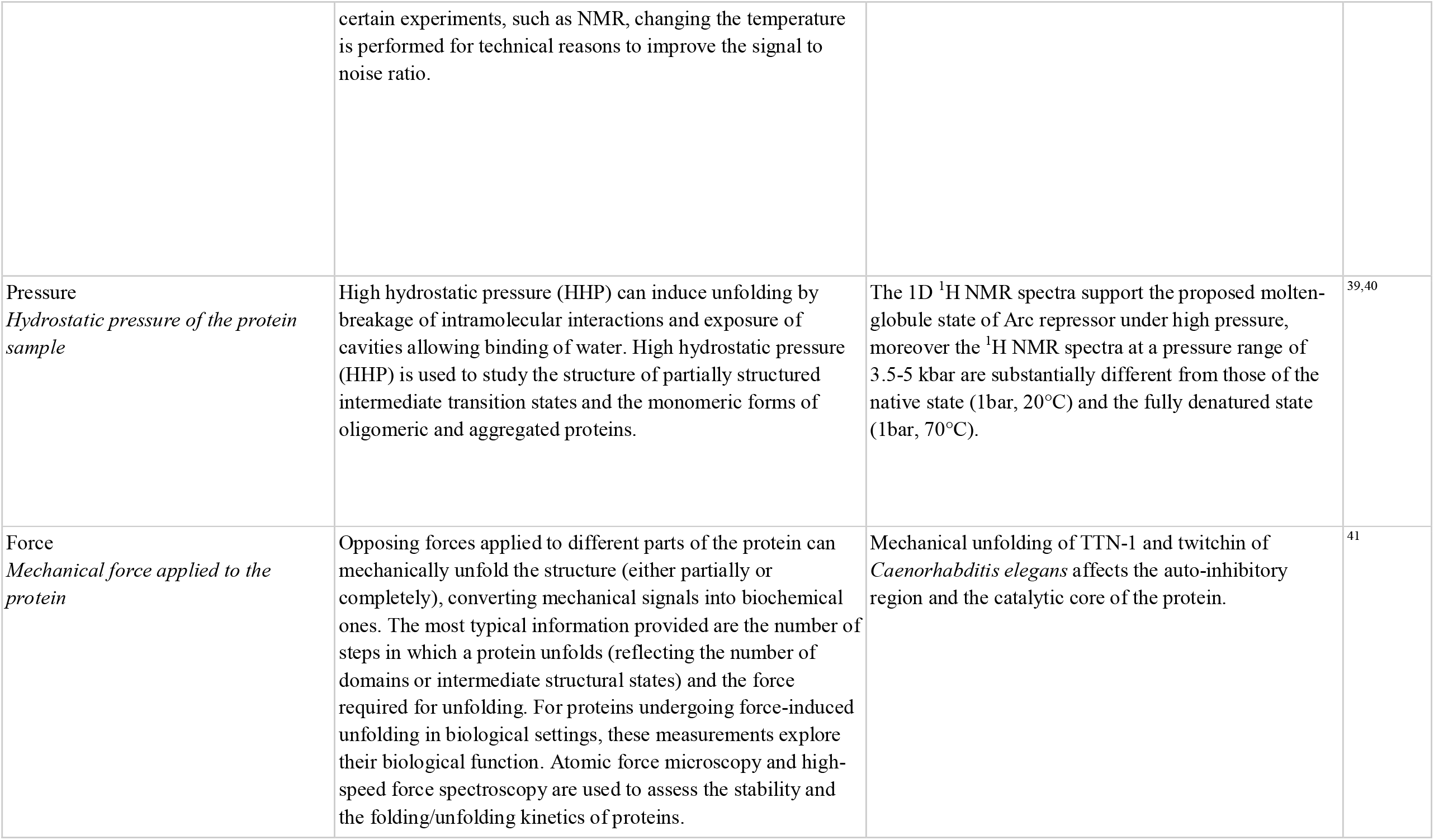
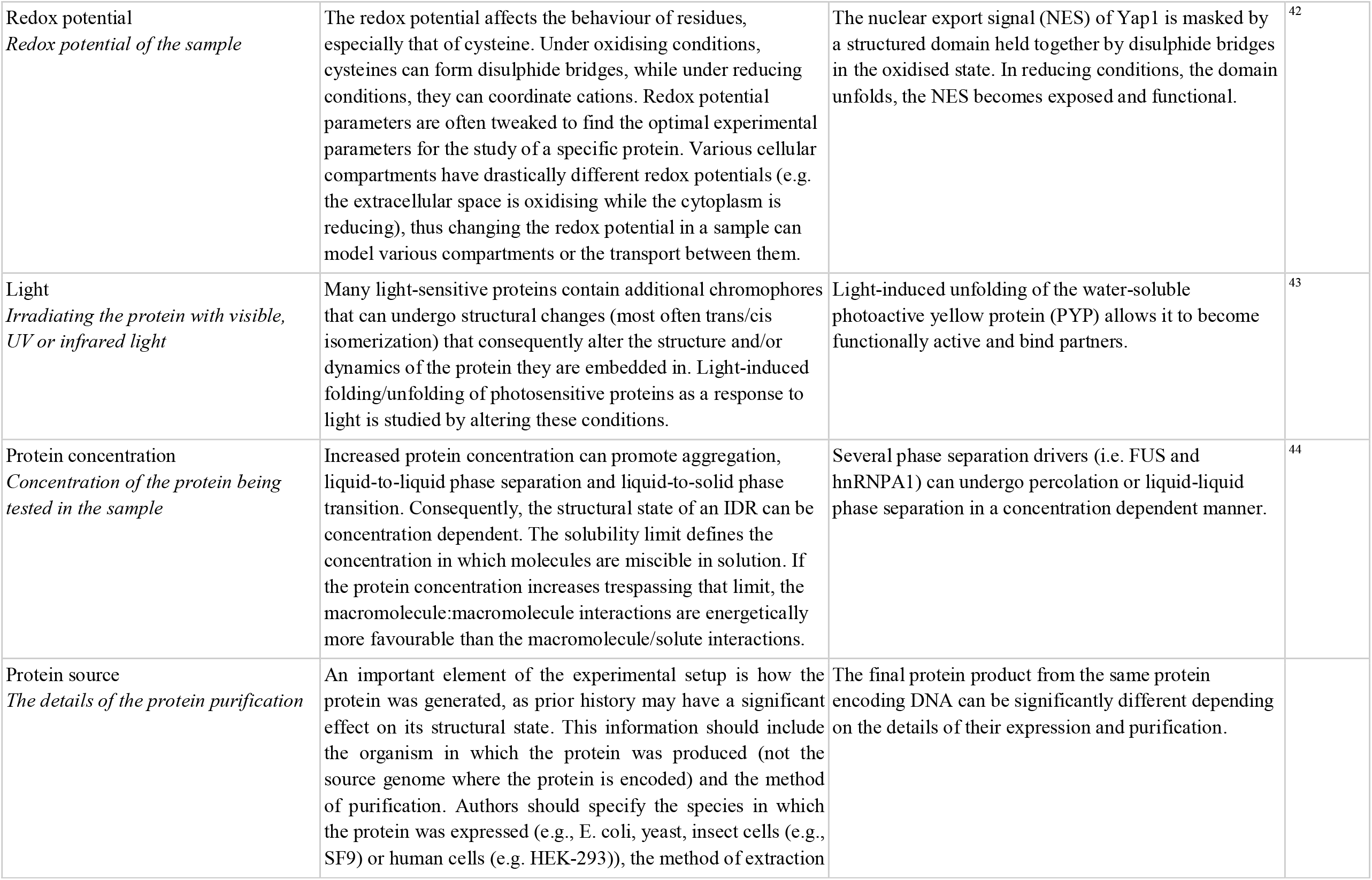
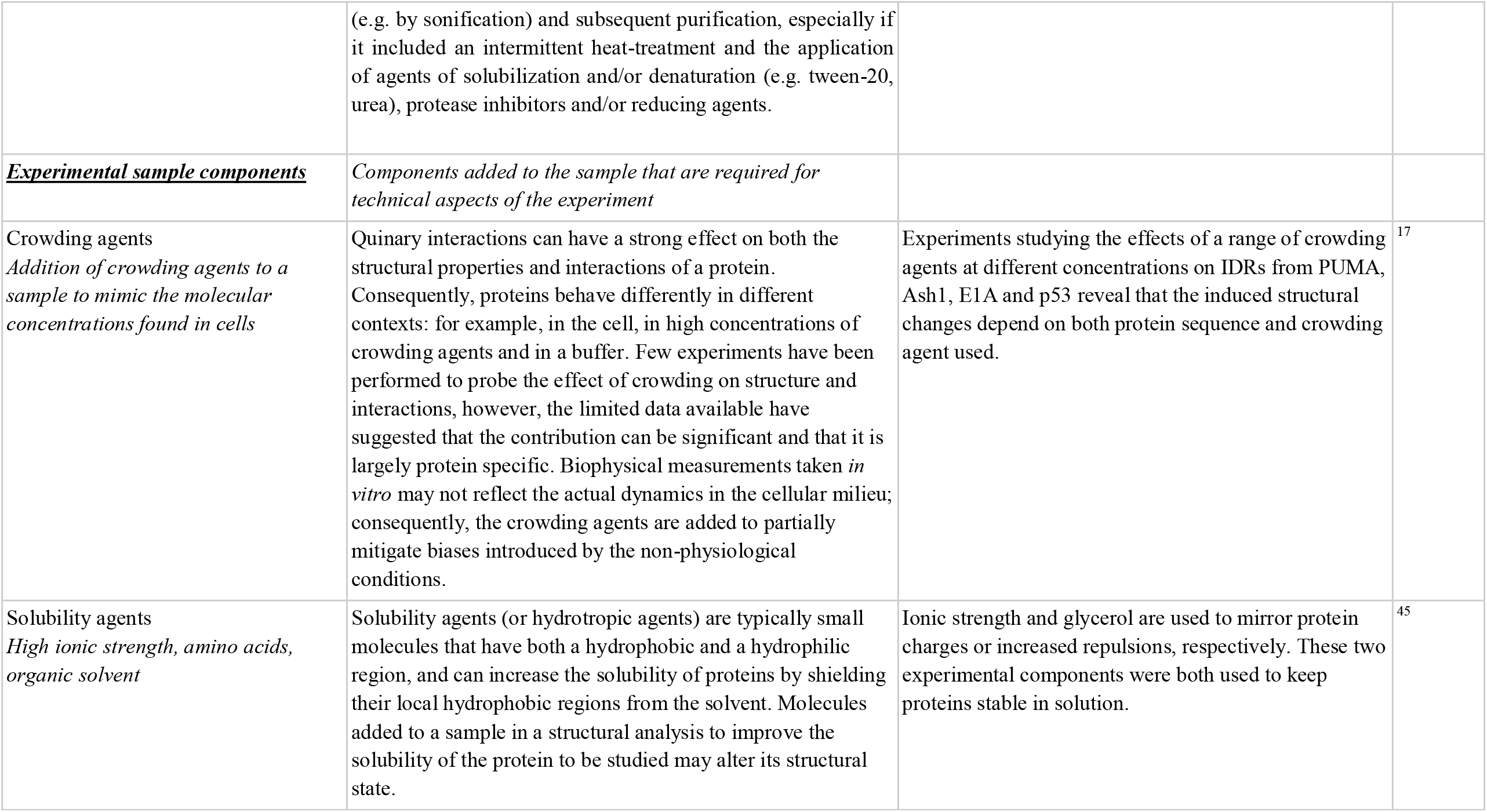
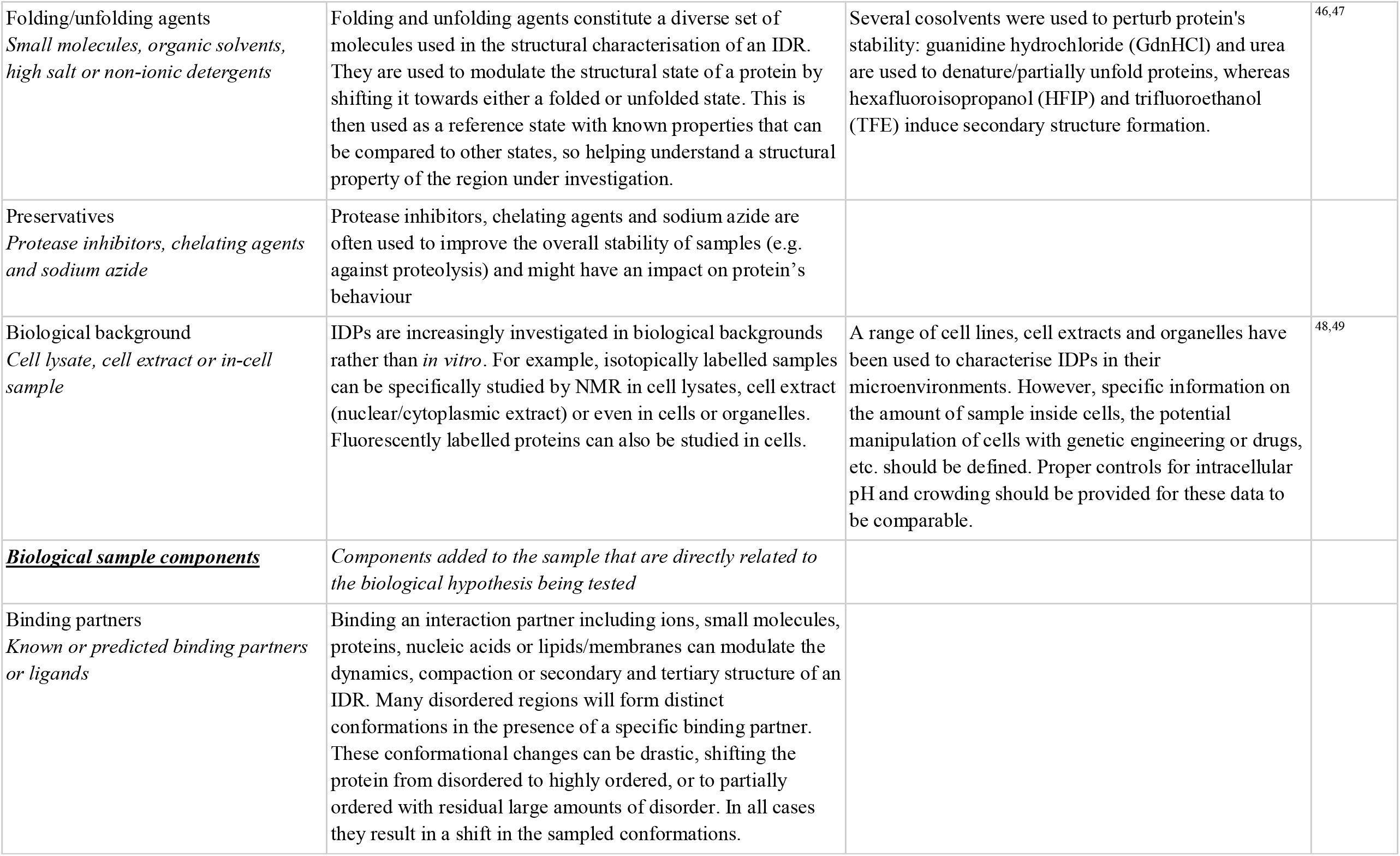
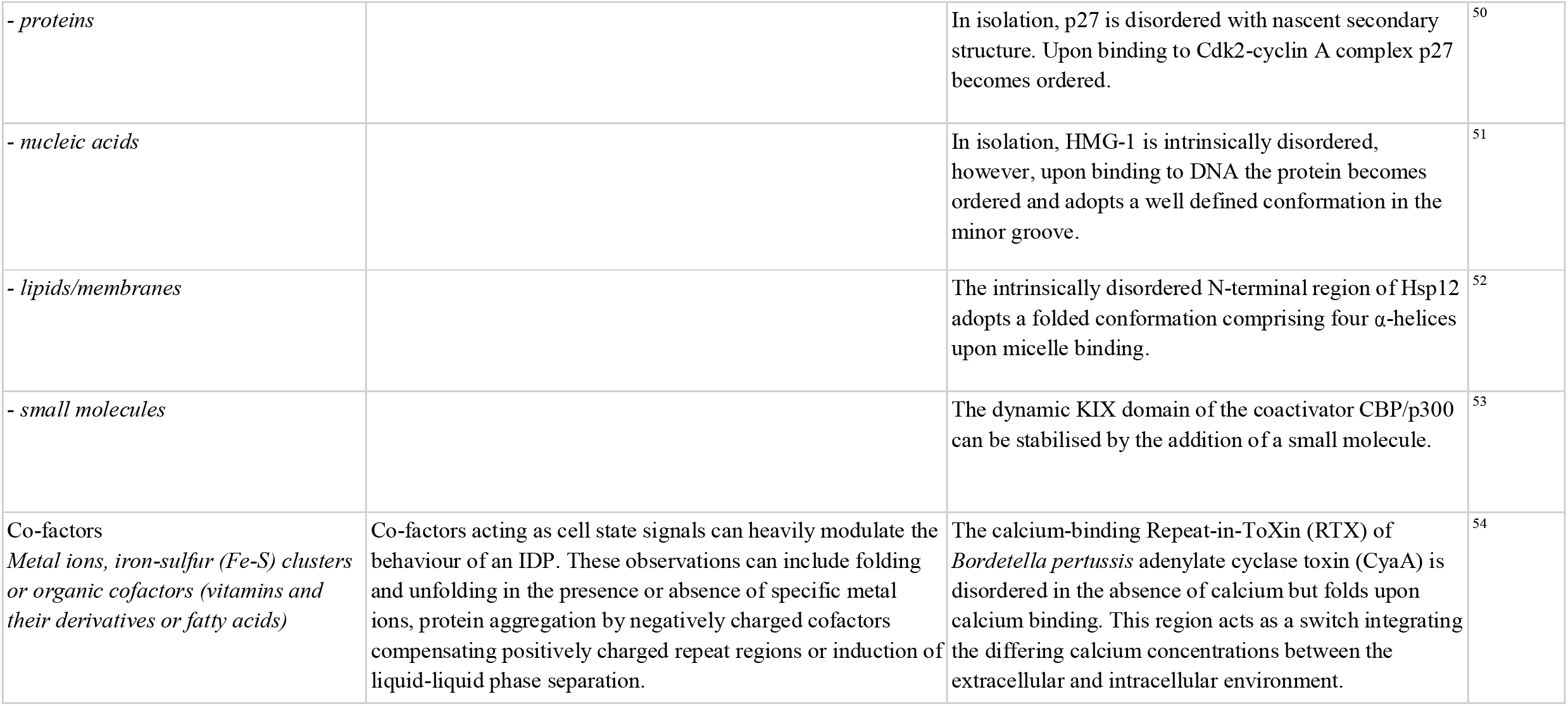
Examples of information that is important to understand the inference that was made from a disorder experiment.
- Any additional components in the sample that could alter the interpretation of the results. This attribute is important to clearly capture structural changes induced by binding partners. However, it also includes other components such as reducing agents, cofactors and crowding agents which may trigger a structural change on the protein of interest. Each component should be defined unambiguously, and if possible, include the concentration of the sample components and refer to external databases including a definition of the molecule (e.g., Uniprot, ChEMBL). Additional protein components should be defined to the same level of detail as the experimental region being studied. See next section and Table 2 for details. *Example: experiment was performed in the presence of 10 g/L polyethylene glycol 400 (PEG400) (CHEMBL:1201478)*.

| Deviation Type | Deviation Description | Example |  |
| --- | --- | --- | --- |
| **<u>Deviations from the canonical protein sequence</u>** | *Definition of the construct used in the experiment* |  |  |
| Post-translational modifications<br>*Covalent modification of a residue side chain* | PTMs can change the physicochemical properties of a sequence and thereby alter the structural state, compaction or dynamics of an IDR. The structures of several IDRs have been shown to be modulated by the addition or removal of a PTM. Studies aimed at investigating these mechanisms will characterise modified proteoforms to understand the structural changes. | 4E-BP2 folds into a 4 stranded beta structure upon phosphorylation of residues T37 and T46. | 8 |
| Substitutions<br>*Replacement of a wild type residue with a non-wild type residue* | Mutations can change the local physicochemical properties of proteins, including charge, hydrophobicity, interaction capacity and size, potentially affecting the stability of folded regions and the solubility of IDRs. Studies introducing mutations can enable the testing of the effect of polymorphisms or disease variants, as well as the effect of certain PTMs (such as phosphomimetics). | A F82K mutation in Ferricytochrome c induces localised unfolding of a distal site in the ferric state. | 31 |
| Tags and labels<br>*Covalent attachment of an entity that enables analysis, identification, purification or solubility of the protein* | Tags and labels can have a measurable influence on the dynamics and stability of the protein they are attached to. The addition of tags is almost always a technical necessity, and its aim is not to measure a biological phenomenon. Tags play three major roles in IDR experiments: (i) for purification (e.g. FLAG tag), (ii) for solubility (e.g. maltose-binding protein (MBP)), (iii) for experimental readout (e.g. fluorescent tags for fluorescent microscopy or paramagnetic tags for NMR). | The addition of a His tag influences myoglobin short time scale (picoseconds) dynamics. | 32 |
| Proteolytic cleavage<br>*Cleavage of the protein chain induced by a protease* | Cleavage can disrupt both local structural elements and long range contacts by increasing the distance between residue pairs. Cleavage also introduces new N- and C-termini in the protein chain changing the polarity, solubility and interaction capacity of regions. Many proteins, especially extracellular proteins, are known to undergo cleavage, often in many subsequent steps. Cleavage products can be created in response to signalling events and often have very different biological activity, interaction capacity and structural states. | Cleavage of the disordered osteopontin removes long range intramolecular interactions, changing the structural state and the accessibility of the integrin binding site. | 33 |
| **<u>Experimental parameters</u>** | *Parameters of the experimental setup for a sample* |  |  |
| pH<br>*pH of the sample* | The pH can affect the strength of ionic and hydrogen bonds and can so modulate the structural state of a protein<sup>34</sup>. Experimental parameters are often tweaked to find the optimal experimental parameters for the study of a specific protein, sometimes resulting in the use of non-physiological pH. Furthermore, comparison of a physiological state with a non-physiological pH state can be used to probe the structural properties of the region of interest. For example, forcing the complete unfolding of a construct with harsh experimental conditions to allow comparison to a "ground state" | NhaA, a sodium proton antiporter of the inner membrane of *Escherichia coli*, is activated at pH values between 6-7, with a maximal activity at pH 8.5, and inactivated by acidic pH. | 35,36 |
| Temperature<br>*Temperature of the sample* | The temperature has an explicit role in determining the strength of entropic terms in the Gibbs free energy that controls the stability of protein structures and complexes. Thus, changing the temperature can drastically change the stability of folded proteins and dynamics of IDRs. Changing the temperature of a protein sample in an experiment can serve to explore its folding/unfolding kinetics, its stability and oligomerization. For calorimetric techniques, such as Differential Scanning Calorimetry (DSC), regulating temperature is what provides the measurable signal. For<br>certain experiments, such as NMR, changing the temperature is performed for technical reasons to improve the signal to noise ratio. | Hp26 becomes active with increased temperature in a two-step mechanism that firstly activates the protein and then unfolds it. | 37,38 |
| Pressure<br>*Hydrostatic pressure of the protein sample* | High hydrostatic pressure (HHP) can induce unfolding by breakage of intramolecular interactions and exposure of cavities allowing binding of water. High hydrostatic pressure (HHP) is used to study the structure of partially structured intermediate transition states and the monomeric forms of oligomeric and aggregated proteins. | The 1D $^1\text{H}$ NMR spectra support the proposed molten-globule state of Arc repressor under high pressure, moreover the $^1\text{H}$ NMR spectra at a pressure range of 3.5-5 kbar are substantially different from those of the native state (1bar, 20°C) and the fully denatured state (1bar, 70°C). | 39,40 |
| Force<br>*Mechanical force applied to the protein* | Opposing forces applied to different parts of the protein can mechanically unfold the structure (either partially or completely), converting mechanical signals into biochemical ones. The most typical information provided are the number of steps in which a protein unfolds (reflecting the number of domains or intermediate structural states) and the force required for unfolding. For proteins undergoing force-induced unfolding in biological settings, these measurements explore their biological function. Atomic force microscopy and high-speed force spectroscopy are used to assess the stability and the folding/unfolding kinetics of proteins. | Mechanical unfolding of TTN-1 and twitchin of *Caenorhabditis elegans* affects the auto-inhibitory region and the catalytic core of the protein. | 41 |
| Redox potential<br>*Redox potential of the sample* | The redox potential affects the behaviour of residues, especially that of cysteine. Under oxidising conditions, cysteines can form disulphide bridges, while under reducing conditions, they can coordinate cations. Redox potential parameters are often tweaked to find the optimal experimental parameters for the study of a specific protein. Various cellular compartments have drastically different redox potentials (e.g. the extracellular space is oxidising while the cytoplasm is reducing), thus changing the redox potential in a sample can model various compartments or the transport between them. | The nuclear export signal (NES) of Yap1 is masked by a structured domain held together by disulphide bridges in the oxidised state. In reducing conditions, the domain unfolds, the NES becomes exposed and functional. | 42 |
| Light<br>*Irradiating the protein with visible, UV or infrared light* | Many light-sensitive proteins contain additional chromophores that can undergo structural changes (most often trans/cis isomerization) that consequently alter the structure and/or dynamics of the protein they are embedded in. Light-induced folding/unfolding of photosensitive proteins as a response to light is studied by altering these conditions. | Light-induced unfolding of the water-soluble photoactive yellow protein (PYP) allows it to become functionally active and bind partners. | 43 |
| Protein concentration<br>*Concentration of the protein being tested in the sample* | Increased protein concentration can promote aggregation, liquid-to-liquid phase separation and liquid-to-solid phase transition. Consequently, the structural state of an IDR can be concentration dependent. The solubility limit defines the concentration in which molecules are miscible in solution. If the protein concentration increases trespassing that limit, the macromolecule:macromolecule interactions are energetically more favourable than the macromolecule/solute interactions. | Several phase separation drivers (i.e. FUS and hnRNPA1) can undergo percolation or liquid-liquid phase separation in a concentration dependent manner. | 44 |
| Protein source<br>*The details of the protein purification* | An important element of the experimental setup is how the protein was generated, as prior history may have a significant effect on its structural state. This information should include the organism in which the protein was produced (not the source genome where the protein is encoded) and the method of purification. Authors should specify the species in which the protein was expressed (e.g., E. coli, yeast, insect cells (e.g., SF9) or human cells (e.g. HEK-293)), the method of extraction (e.g. by sonification) and subsequent purification, especially if it included an intermittent heat-treatment and the application of agents of solubilization and/or denaturation (e.g. tween-20, urea), protease inhibitors and/or reducing agents. | The final protein product from the same protein encoding DNA can be significantly different depending on the details of their expression and purification. |  |
| **<u>Experimental sample components</u>** | *Components added to the sample that are required for technical aspects of the experiment* |  |  |
| Crowding agents<br>*Addition of crowding agents to a sample to mimic the molecular concentrations found in cells* | Quinary interactions can have a strong effect on both the structural properties and interactions of a protein. Consequently, proteins behave differently in different contexts: for example, in the cell, in high concentrations of crowding agents and in a buffer. Few experiments have been performed to probe the effect of crowding on structure and interactions, however, the limited data available have suggested that the contribution can be significant and that it is largely protein specific. Biophysical measurements taken *in vitro* may not reflect the actual dynamics in the cellular milieu; consequently, the crowding agents are added to partially mitigate biases introduced by the non-physiological conditions. | Experiments studying the effects of a range of crowding agents at different concentrations on IDRs from PUMA, Ash1, E1A and p53 reveal that the induced structural changes depend on both protein sequence and crowding agent used. | <sup>17</sup> |
| Solubility agents<br>*High ionic strength, amino acids, organic solvent* | Solubility agents (or hydrotropic agents) are typically small molecules that have both a hydrophobic and a hydrophilic region, and can increase the solubility of proteins by shielding their local hydrophobic regions from the solvent. Molecules added to a sample in a structural analysis to improve the solubility of the protein to be studied may alter its structural state. | Ionic strength and glycerol are used to mirror protein charges or increased repulsions, respectively. These two experimental components were both used to keep proteins stable in solution. | <sup>45</sup> |
| Folding/unfolding agents<br>*Small molecules, organic solvents, high salt or non-ionic detergents* | Folding and unfolding agents constitute a diverse set of molecules used in the structural characterisation of an IDR. They are used to modulate the structural state of a protein by shifting it towards either a folded or unfolded state. This is then used as a reference state with known properties that can be compared to other states, so helping understand a structural property of the region under investigation. | Several cosolvents were used to perturb protein's stability: guanidine hydrochloride (GdnHCl) and urea are used to denature/partially unfold proteins, whereas hexafluoroisopropanol (HFIP) and trifluoroethanol (TFE) induce secondary structure formation. | 46,47 |
| Preservatives<br>*Protease inhibitors, chelating agents and sodium azide* | Protease inhibitors, chelating agents and sodium azide are often used to improve the overall stability of samples (e.g. against proteolysis) and might have an impact on protein's behaviour |  |  |
| Biological background<br>*Cell lysate, cell extract or in-cell sample* | IDPs are increasingly investigated in biological backgrounds rather than *in vitro*. For example, isotopically labelled samples can be specifically studied by NMR in cell lysates, cell extract (nuclear/cytoplasmic extract) or even in cells or organelles. Fluorescently labelled proteins can also be studied in cells. | A range of cell lines, cell extracts and organelles have been used to characterise IDPs in their microenvironments. However, specific information on the amount of sample inside cells, the potential manipulation of cells with genetic engineering or drugs, etc. should be defined. Proper controls for intracellular pH and crowding should be provided for these data to be comparable. | 48,49 |
| **<u>Biological sample components</u>** | *Components added to the sample that are directly related to the biological hypothesis being tested* |  |  |
| Binding partners<br>*Known or predicted binding partners or ligands* | Binding an interaction partner including ions, small molecules, proteins, nucleic acids or lipids/membranes can modulate the dynamics, compaction or secondary and tertiary structure of an IDR. Many disordered regions will form distinct conformations in the presence of a specific binding partner. These conformational changes can be drastic, shifting the protein from disordered to highly ordered, or to partially ordered with residual large amounts of disorder. In all cases they result in a shift in the sampled conformations. |  |  |
| - *proteins* |  | In isolation, p27 is disordered with nascent secondary structure. Upon binding to Cdk2-cyclin A complex p27 becomes ordered. | 50 |
| - *nucleic acids* |  | In isolation, HMG-1 is intrinsically disordered, however, upon binding to DNA the protein becomes ordered and adopts a well defined conformation in the minor groove. | 51 |
| - *lipids/membranes* | | The intrinsically disordered N-terminal region of Hsp12 adopts a folded conformation comprising four α-helices upon micelle binding. | 52 |
| - *small molecules* |  | The dynamic KIX domain of the coactivator CBP/p300 can be stabilised by the addition of a small molecule. | 53 |
| Co-factors<br>*Metal ions, iron-sulfur (Fe-S) clusters or organic cofactors (vitamins and their derivatives or fatty acids)* | Co-factors acting as cell state signals can heavily modulate the behaviour of an IDP. These observations can include folding and unfolding in the presence or absence of specific metal ions, protein aggregation by negatively charged cofactors compensating positively charged repeat regions or induction of liquid-liquid phase separation. | The calcium-binding Repeat-in-ToXin (RTX) of *Bordetella pertussis* adenylate cyclase toxin (CyaA) is disordered in the absence of calcium but folds upon calcium binding. This region acts as a switch integrating the differing calcium concentrations between the extracellular and intracellular environment. | 54 |

In the case of data being stored in a database, transferred between resources, or defined in the absence of a paper, it is important to also include the source of the data.

##### Data source

a reference to where the data were originally described.

- In cases where data were published in a paper, the following information should be provided:
  - publication database and identifier *Example: PMID: 35055108*
- In cases where data were directly submitted to a data resource, the following information should be provided:
  - the name of the data resource
  - the accession number of the record holding the data in that resource
  - the data creator who submitted the data
  - contact details of the data creator

### Key factors that can influence the inference made from a disorder experiment

Numerous factors connected to the protein region, protein construct or the experimental setup can influence the structural state of the protein region being studied and, consequently, our confidence in the biological relevance of the observed structure (see Table 2)^17,18^. These factors can be experimental perturbations, to allow experimental measurements to be collected, or biological perturbations, to understand biologically relevant proteoforms, conditions or samples. In these cases, any description of the structural state is only meaningful when the relevant factors that influence the observed state are specified. While the minimum information requires the protein region and the experimental method to be defined, it is up to the discretion of the authors to report deviations from the established protocol, sample or sequence that could alter the interpretation of the results. Consequently, an explicit statement by an author will simplify the task of the curator or reader to make a judgement of the importance of a given deviation. In complex cases the meaningful description of the inferred structural states can include several pieces of information that go beyond the specification of the protein region and the experimental method applied. In Table 2, we provide pointers on which factors might be considered important deviations based on known biological cases of conditional protein disorder and common experimental perturbations.

### Example use cases

There are several use cases for MIADE (Table 1), however, in practice there are two major distinct applications: (i) creating an unambiguous description of an experiment in free text and (ii) encoding the fundamental unit of metadata for an experiment in a standardised format. In this section, we will give examples of how MIADE can be applied in each of these cases.

#### MIADE for authors

A key step in data capture is the unambiguous description of the expert interpretation of the primary data. Consequently, an accurate and unequivocal definition of the experimental observation in the text of an article that adheres to the MIADE guidelines will simplify all downstream data interpretation. Defining an experiment in free text requires detail that allows the experiment to be fully reproduced. Consequently, most articles describe the experimental detail at a level of granularity that far exceeds the requirements of a MIADE compliant entry. However, a comprehensive description of an experiment’s design and results does not mean that the data is accessible to the wider biological community. A common issue amongst non-expert readers and curators is that the data is described in a manner that is highly technical, requires extensive knowledge of the experimental method or uses field-specific jargon. Furthermore, important details are often not apparent as they are in materials and methods sections, supplementary materials or even a previously published paper. Consequently, MIADE guidelines recommend an explicit and unambiguous description of the experimental design, the proteins under analysis and the expert interpretation of the results.

Consideration should be given to the fact that the description should be understandable to non-experts in the wider biological community and the key data should be explicitly stated. This will improve the clarity of the document and allow rapid annotation by curators for community resources. In many cases, writing engaging and readable scientific prose, and writing unequivocal descriptions of complex experiments are conflicting goals. However, in any case where such conflicts occur, substance should take precedence over style. For example, the definition of a protein as *“Budding Yeast (Saccharomyces cerevisiae strain ATCC 204508 / S288c (TaxID:559292)) Spindle assembly checkpoint component MAD3 (MAD3) (UniProt:P47074)”* may be rather awkward when compared to*”yeast MAD3”*. However, it removes ambiguity from the protein definition. By following the examples in the checklist and understanding that a reader may not be an expert, data can be presented in a manner that is both accurate and globally accessible.

#### MIADE implementation in DisProt

An important aspect to represent experimentally determined structural states of IDPs and IDRs in a standard format is the use of stable external identifiers and controlled vocabularies (CV) to unambiguously describe the captured data. In the future, IDP-specific exchange formats should be developed to define these attributes for experimental metadata, however, for the moment it is useful to consider how DisProt stores MIADE compliant data.

DisProt is a manually curated resource of intrinsically disordered regions (IDRs) and proteins (IDPs) from literature, that relies on both professional and community curation. All DisProt entries correspond to a specific UniProt entry (or one of its isoforms) and describe the structural state(s) of the region(s) of the protein. When available, information on the presence of transitions between states, interactions and functions, is also curated. The annotation of structural states and transitions makes use of specific IDPO terms (https://disprot.org/ontology). As part of the development of MIADE guidelines we have updated the DisProt database and curation framework to allow the annotation of MIADE-compliant entries^1^. An improved construct definition was required to encode tags, labels, mutations or modifications and the experimental setup definition was updated to allow complex experimental samples to be described. Importantly, these additions will allow DisProt to annotate the observations of complex experiments defining conditional multistate IDRs that are becoming increasingly common in the literature.

##### Proteoform definition

The DisProt resource already included an unambiguous definition of the protein or protein isoforms (using UniProt accession numbers) and its regions by mapping to the UniProt sequence. The updated implementation can now define non-canonical and modified proteoforms. The MIADE integration allows the possibility to encode deviations from the wildtype UniProt defined protein sequence. Furthermore, the complete sequence of the experimental construct can now be annotated if available. Annotatable construct alterations include *tags* and *labels* (using the PSI-MI ontology (https://www.ebi.ac.uk/ols/ontologies/mod)^19^), *mutations* (using the HGSV nomenclature (https://varnomen.hgvs.org/)) and *PTMs* and *non-standard amino acids* (using the PSI-MOD ontology (https://www.ebi.ac.uk/ols/ontologies/mod)^20^).

##### Experimental conditions definition

DisProt uses the Evidence and Conclusion Ontology (ECO, http://www.evidenceontology.org/)^21^ to annotate experimental methods. In addition, the DisProt database can now store a range of experimental parameters that can influence our understanding of the biological relevance of an experimental observation, i.e. *pH, temperature, pressure, ionic strength*, and *oxidation-reduction potential*. The parameter can be quantified in cases where this information is available. All parameters are defined in the NCI Thesaurus OBO Edition controlled vocabulary (https://ncit.nci.nih.gov/ncitbrowser/) and their units in the Units of Measurement Ontology (https://bioportal.bioontology.org/ontologies/UO). Deviations from the expected value in the experiment parameter (e.g. within normal range, increased, decreased, not specified or not relevant) can also be added. All information is annotated with the text description taken directly from the scientific article and curators’ statements can be added to further clarify annotation.

##### Experimental components definition

The DisProt database can now describe experimental sample components such as lipids, nucleic acids, small molecules, metal ions or proteins present during the characterisation of the structural state of an IDR. The concentration of the components and cross reference to the specific database, i.e. CheBI^22^, ENA^23^, RNAcentral^24^ and UniProt^13^ can also be added. Similar to the other MIADE fields, a text description can be added into the corresponding *Statement* field.

A representative list of DisProt use cases highlighting novel information covered by the addition of fields from the MIADE update is described in Table 3.

**Table 3.**
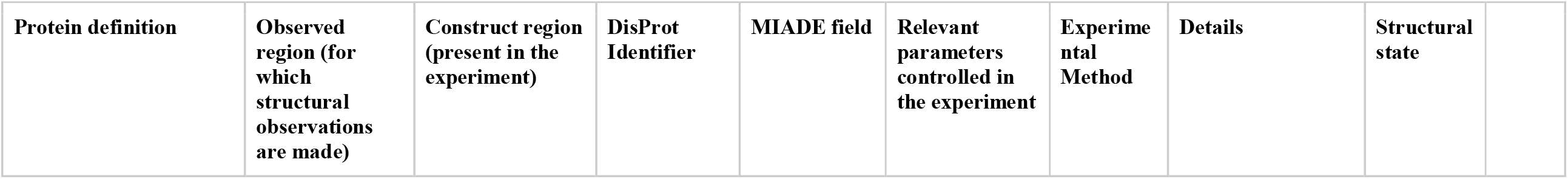
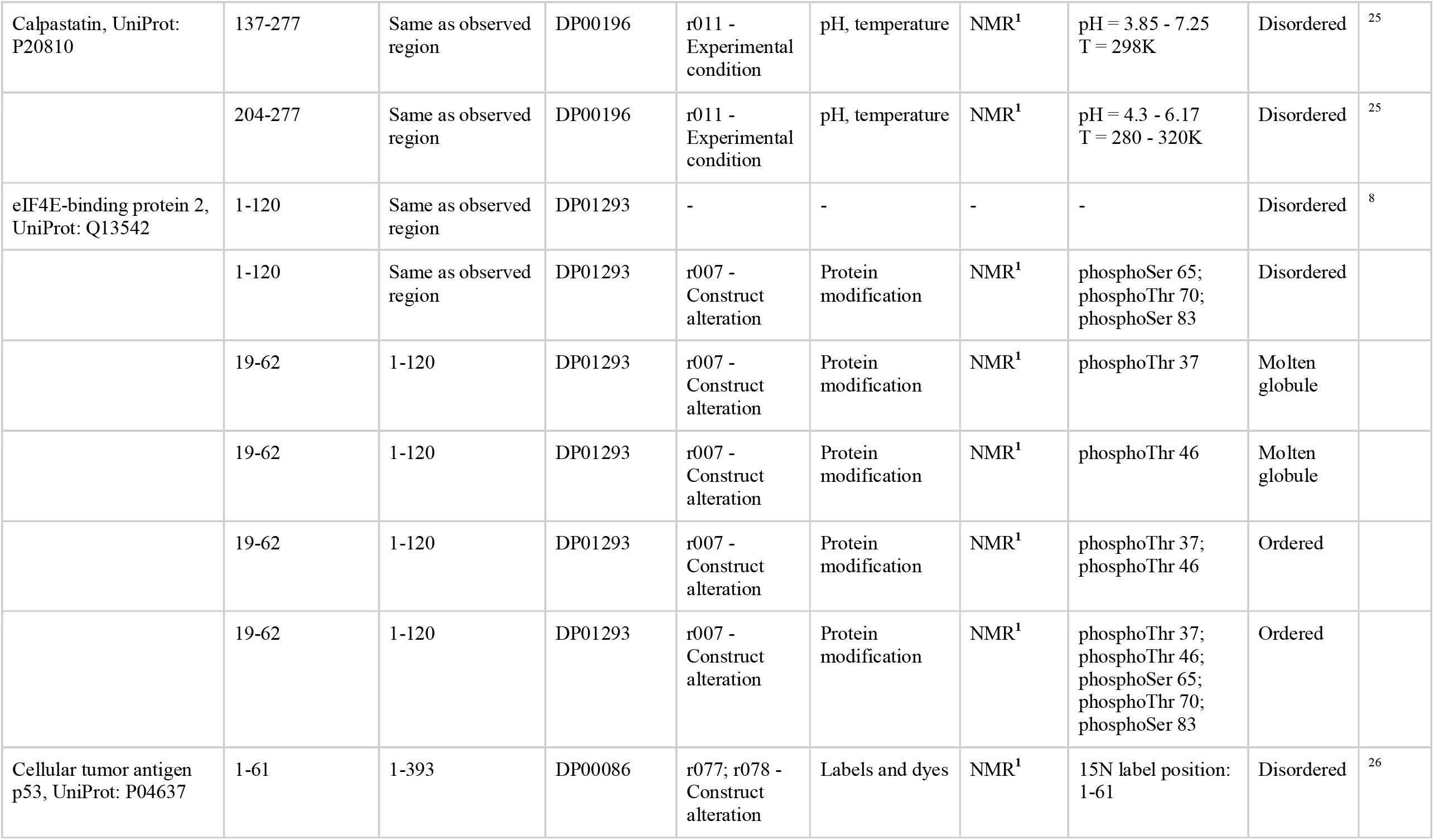
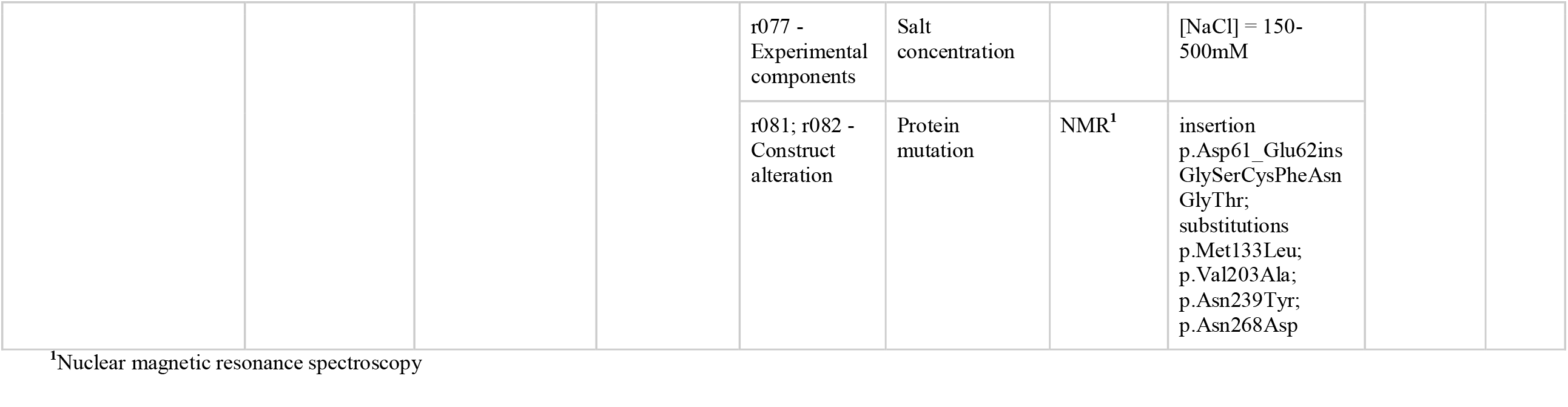
Extra data curated by DisProt to allow a MIADE compliant annotation for the case study examples.

| Protein definition | Observed region (for which structural observations are made) | Construct region (present in the experiment) | DisProt Identifier | MIADE field | Relevant parameters controlled in the experiment | Experimental Method | Details | Structural state |
| --- | --- | --- | --- | --- | --- | --- | --- | --- |

|  |  |  |  |  |  |  |  |  |  |
| --- | --- | --- | --- | --- | --- | --- | --- | --- | --- |
| Calpastatin, UniProt: P20810 | 137-277 | Same as observed region | DP00196 | r011 - Experimental condition | pH, temperature | NMR<sup>1</sup> | pH = 3.85 - 7.25<br>T = 298K | Disordered | <sup>25</sup> |
|  | 204-277 | Same as observed region | DP00196 | r011 - Experimental condition | pH, temperature | NMR<sup>1</sup> | pH = 4.3 - 6.17<br>T = 280 - 320K | Disordered | <sup>25</sup> |
| eIF4E-binding protein 2, UniProt: Q13542 | 1-120 | Same as observed region | DP01293 | - | - | - | - | Disordered | <sup>8</sup> |
|  | 1-120 | Same as observed region | DP01293 | r007 - Construct alteration | Protein modification | NMR<sup>1</sup> | phosphoSer 65;<br>phosphoThr 70;<br>phosphoSer 83 | Disordered |  |
|  | 19-62 | 1-120 | DP01293 | r007 - Construct alteration | Protein modification | NMR<sup>1</sup> | phosphoThr 37 | Molten globule |  |
|  | 19-62 | 1-120 | DP01293 | r007 - Construct alteration | Protein modification | NMR<sup>1</sup> | phosphoThr 46 | Molten globule |  |
|  | 19-62 | 1-120 | DP01293 | r007 - Construct alteration | Protein modification | NMR<sup>1</sup> | phosphoThr 37;<br>phosphoThr 46 | Ordered |  |
|  | 19-62 | 1-120 | DP01293 | r007 - Construct alteration | Protein modification | NMR<sup>1</sup> | phosphoThr 37;<br>phosphoThr 46;<br>phosphoSer 65;<br>phosphoThr 70;<br>phosphoSer 83 | Ordered |  |
| Cellular tumor antigen p53, UniProt: P04637 | 1-61 | 1-393 | DP00086 | r077; r078 - Construct alteration | Labels and dyes | NMR<sup>1</sup> | <sup>15</sup>N label position: 1-61 | Disordered | <sup>26</sup> |
|  |  |  |  | r077 -<br>Experimental<br>components | Salt<br>concentration |  | [NaCl] = 150-500mM |  |
|  |  |  |  | r081; r082 -<br>Construct<br>alteration | Protein<br>mutation | NMR<sup>1</sup> | insertion<br>p.Asp61_Glu62ins<br>GlySerCysPheAsn<br>GlyThr;<br>substitutions<br>p.Met133Leu;<br>p.Val203Ala;<br>p.Asn239Tyr;<br>p.Asn268Asp |  |

<sup>1</sup>Nuclear magnetic resonance spectroscopy

### Case studies

While MIADE only captures the core structural inferences derived from disorder experiments, it can be applied to the description of experimental data with a very wide range of complexity in terms of experimental design and studied system. In the following section we demonstrate how MIADE-compliant information can be created using as examples three papers applying nuclear magnetic resonance (NMR) in increasingly complex setups. These extracts serve as examples of good practice and are accompanied by a MIADE compliant entry in the DisProt resource (Table 3). We highlight the three key areas covered by MIADE from each paper: the definition of the protein construct used; the deviation from the wildtype proteoform (including mutations, post-translational modifications, tags, labels and dyes); and the definition of the experimental setup, including the environmental conditions and sample compositions that might have relevance for the structural state.

The first paper describes the disordered structural state of human calpastatin (CAST), an inhibitor of calpain, the Ca^2+^ activated cysteine protease^25^. The authors unambiguously define two protein constructs they used by referencing the common name of the protein and source organism, together with a UniProt accession (‘^*15*^*N-labeled and* ^*13*^*C-labeled full-length hCSD1 [corresponding to A137-K277 of human calpastatin, SwissProt entry P20810]*’ and ‘*C-terminal half of calpastatin (position in whole calpastatin P204-K277)*’). In addition, they also clearly define the experimental method as various forms of NMR experiments, including heteronuclear single quantum coherence (HSQC), calculation of the secondary chemical shift and ^3^J_HNHα_ scalar coupling constants determined with 3D HNCA-E.COSY type experiments. For these experiments, the relevant environmental conditions are temperature and pH, which the authors define in the materials and methods sections (‘*HSQC spectra collected at 298 K and at pH 4*.*3, 5*.*23, and 6*.*17 for hCSD1(67-141) as well as pH 3*.*85, 5*.*53, 6*.*07, and 7*.*25 for hCSD1. The temperature dependence of the same type of resonances was measured at 280, 300, and 320 K in aqueous solution for hCSD1(67-141)*’). Using these setups, the authors then determine that both constructs are essentially disordered and that this observation is largely independent of temperature and pH in the ranges explored. This information together constitute what MIADE can capture, albeit there are more refined observations about the structural properties of the protein, such as: ‘*subdomains A and B, two characteristic binding and functional sites of the inhibitor, have some helical character*’ or ‘*restricted motions on a subnanosecond time scale indicated by larger than average J(0) values are observed for G13-M17, K68-L72, S101-C105, and S128-V132. These residues of restricted mobility also present some residual local structural features highlighted both by secondary chemical shifts, SCS, and by their hydrophobicity pattern*’.

The second paper details disorder experiments performed on Eukaryotic translation initiation factor 4E-binding protein 2 (EIF4EBP2), an interacting partner of Eukaryotic translation initiation factor 4E (eIF4E)^8^. The authors define the protein construct as the full-length human protein by referencing its common name (4E-BP2). The HUGO Gene Nomenclature Committee (HGNC) gene name is EIF4EBP2, and no unambiguous identifier is provided, however, the naming is specific enough to unambiguously identify the protein being studied, given that the protein has no known alternative isoforms. In addition, throughout the paper the authors reference several key residues in the protein (such as T37, T46, S65, T70 and S83) based on which readers and curators can confirm whether they map to the correct UniProt sequence. As opposed to the previous example where conditions were changed, in this case, measurements were performed on distinct proteoforms of the protein. The main structural conclusion of the paper is that the structural state of EIF4EBP2 is dependent on its phosphorylation state. HSQC NMR spectrum shows that ‘*non-phosphorylated 4E-BP2 has intense peaks with narrow* ^*1*^*HN chemical shift dispersion characteristic of IDPs […] However, wild-type 4E-BP2 uniformly phosphorylated at T37, T46, S65, T70 and S83 shows widespread downfield and upfield chemical shifts for residues spanning T19–R62, suggesting folding upon phosphorylation*’. Using partial phosphorylation, the authors then disentangle the individual contribution of each phosphorylation to the induced folding, stating: ‘*No significant change in global dispersion was observed for 4E-BP2 phosphorylated only at S65/T70/S83, demonstrating that it remains disordered, while phosphorylating T37 and T46 (pT37pT46) induces a 4E-BP2 fold identical to phosphorylated wild type. Interestingly, when phosphorylated individually, pT37 or pT46 result in a partly folded state, with some chemical shift changes indicative of ordered structure (pT37). […] Thus, phosphorylation of both T37 and T46 is necessary and sufficient for phosphorylation-induced folding of 4E-BP2*’. The authors also measure the structural effect of binding to eIF4E and find that the interaction induces partial folding of the phosphorylated 4E-BP2: ‘*The spectrum of pT37pT46 in isolated and eIF4E-bound states demonstrate an order-to-disorder transition upon eIF4E binding. […] pT37pT46 undergoes an order-to-disorder transition upon binding to eIF4E*’. Therefore, both phosphorylation and the presence of a binding partner can induce a structural transition of EIF4EBP2 through different mechanisms, and therefore the inference that EIF4EBP2 is disordered is dependent on the exact proteoform as well as the presence of other proteins. In addition to the structural state, the authors also directly address the connection between phosphorylation and the interaction capacity: ‘*non-phosphorylated or minimally phosphorylated 4E-BPs interact tightly with eIF4E, while the binding of highly phosphorylated 4E-BPs is much weaker and can be outcompeted by eIF4G*’. While this piece of information is key to understanding the biological regulatory role of EIF4EBP2, it cannot be captured in the structural state-focused framework of MIADE and should be encoded as additional information in databases.

In the third example, the authors study the human Cellular tumour antigen p53 (TP53) focusing on the structural features of the disordered N-terminal region^26^. The authors clearly define the protein being studied by stating it is human TP53. In addition, they also provide an overview figure that contains the UniProt region boundaries of various p53 regions and domains that are used in the constructs. In contrast to the previous examples, the main construct used in this study is not a full-length protein or an isolated protein region, but a chimeric protein consisting of an isotopically labelled N-terminal and a non-labelled C-terminal region. The authors use a split intein splicing to produce the isotopically labelled disordered N-terminal region and then splice it together with the unlabelled central and C-terminal regions (‘*we utilized intein splicing to segmentally label the NTAD within tetrameric p53 [*…*] NTAD (residues 1–61) labeled with an NMR-active isotope (*^*15*^*N), while residues 62–393 remained unlabeled and NMR invisible*’). As a result of this technique, the final construct has a short insertion where the intein was located, the position of which was carefully chosen: ‘*The intein splice site was selected as D61/E62, a site that is distant in the amino acid sequence from interaction sites or well-folded domains. Careful selection of the splice site is important, since the Npu DnaE intein system inserts nonnative residues (GSCFNGT in the p53 constructs used here) at the splice site*.’ This construct enables the assessment of the structural state of the disordered NTAD in the context of the full length tetrameric TP53 by NMR HSQC spectra. For technical reasons, the authors further introduced mutations to the sequence outside the disordered regions being studied: ‘*To improve expression levels, stabilizing mutations (M133L/V203A/N239Y/N268D) were introduced into the DNA-binding domain*’. The definition of the environmental conditions covers the temperature and salt concentrations, with all other parameters supposedly being in the normal range of similar NMR measurements: ‘*unless otherwise stated, all spectra were recorded at 25 °C for samples in NMR buffer*’ and ‘*salt titrations for p53(1–312) and p53(1–61) were carried out with protein concentrations of 150* μ*M. The initial titration point had a NaCl concentration of 150 mM, and NaCl from a 5-M concentrated stock was added to this sample at 50-mM increments up to 500 mM NaCl*’. Apart from unambiguously defining the protein construct, the proteoform, the techniques and the environmental conditions, the main conclusion about the structural state is also clearly stated as: ‘*the HSQC spectrum of the NTAD-p53 tetramer shows that the NTAD remains dynamically disordered in the full-length protein*’. Like the previous papers, this work contains a considerable number of non-structural observations. These include data on the interaction between the folded DNA-binding domain (DBD), the disordered NTAD and cognate and non-cognate DNA - additions to the experimental sample in certain experiments, the sequences of which are also defined. While this information is outside of the scope of MIADE, it is again written in a clear way enabling specific databases that go beyond MIADE to capture these observations in a structured way.

### Towards standards for the complete definition of an IDP experiment

The MIADE guidelines provide simple recommendations for the definition of the minimal useful level of experimental metadata. However, they are just the first step towards the standardisation and several additional developments are required to standardise the IDP field.

#### Standardised exchange format

The MIADE guidelines do not define a structured data format. The IDP field requires exchange formats that can hold experimental data at a range of detail from a MIADE-compliant definition to a description of the experiment and results that would allow the experiment to be recreated (Figure 1B). Furthermore, they should include the possibility to store raw and processed experimental measurements in addition to interpreted structural observations derived from the data. Given the heterogeneity of the methodology applied to the field of IDP research there is also a strong requirement for experimental method specific standardised exchange formats. These developments should be driven by the existing communities and resources for these methods, however, given the parallel requirements across many of these fields, efforts should be made to collaborate and reuse structured data formats when possible. The development of exchange formats and associated controlled vocabularies to permit the dissemination and storage of data relating to IDR structure and function has begun under the HUPO Proteomics Standards Initiative funded by ELIXIR.

#### Controlled vocabularies and ontologies

Controlled vocabularies and ontologies are a key component of the standardisation of IDP data storage. These definitions standardise the meaning of the terms used to describe the IDP data allowing the complete unambiguous annotation of an IDP experiment and results. Recent additions to the Intrinsically Disordered Proteins Ontology (IDPO) and the Evidence and Conclusion Ontology (ECO) have significantly increased the coverage of IDP-related terms. Yet, both ontologies are still incomplete and additional terms are necessary including those related to newly developed experimental approaches, computational methods, non-binary structural classifications (i.e., more detailed than order/disorder including dynamics, secondary structure propensity and compaction), structural transition definitions and conditionality. However, ongoing work by the DisProt community is continually adding terms as required.

#### Suitability for past, present and future

Any standardised exchange format, controlled vocabularies and ontologies developed over the coming years will require flexibility to hold the heterogeneous data that have been produced over the past decades. Furthermore, they will require significant and ongoing development to cover information from the ever-improving methodology being developed by the IDP community. Consequently, we see the standardisation of the IDP fields as a long-term project requiring extensive consultation between the experimental, computational and database communities within the IDP field. Areas where recent advances have outpaced the resources that store the available data are residue centric information, descriptions of dynamics and conditionality, and structural observations beyond the classical binary order/disorder classification. The detailed definition of the structural states, state conditions and inter-state transitions of a region is key to capturing these data and should be at the core of any standard development.

### MIADE-compliant metadata capture at source

To date, direct submission of data to community resources is underutilised by the IDP community. IDP resources should improve their capacity to receive data pre-publication including the possibility to embargo data until the time of final publication (similar to the PDB model) and develop tools and resources that simplify MIADE-compliant reporting. Furthermore, the IDP community should enforce the deposition of experimental data and metadata as a required component of the publication process. The ideal situation would include the pre-publication submission of primary source data directly to the corresponding field specific resource (e.g. Biological Magnetic Resonance Bank (BMRB) for NMR data^27^, Protein Circular Dichroism Data Bank (PCDDB) for circular dichroism data^28^, SASBDB for Small-Angle X-ray Scattering (SAXS) data^29^ or Protein Ensemble Database (PED) for protein ensemble data^30^). Subsequently, a reference to primary source data and MIADE-compliant experimental metadata should then be submitted to a community resource such as DisProt or IDEAL^1,2^. This benefits the databases, as the efficiency of data collection and verification is increased. This in turn benefits the IDP community and wider biological community, as more and more precise data, linked to related primary data in field-specific databases, are readily available. Currently, several databases allow pre-or post-publication submission of data related to IDR experiments, each with their own submission process and data formats (Table 4). However, the proportion of data created that is captured by these resources varies widely and no resource is successful in capturing all data produced that fall within their scope.

**Table 4.**
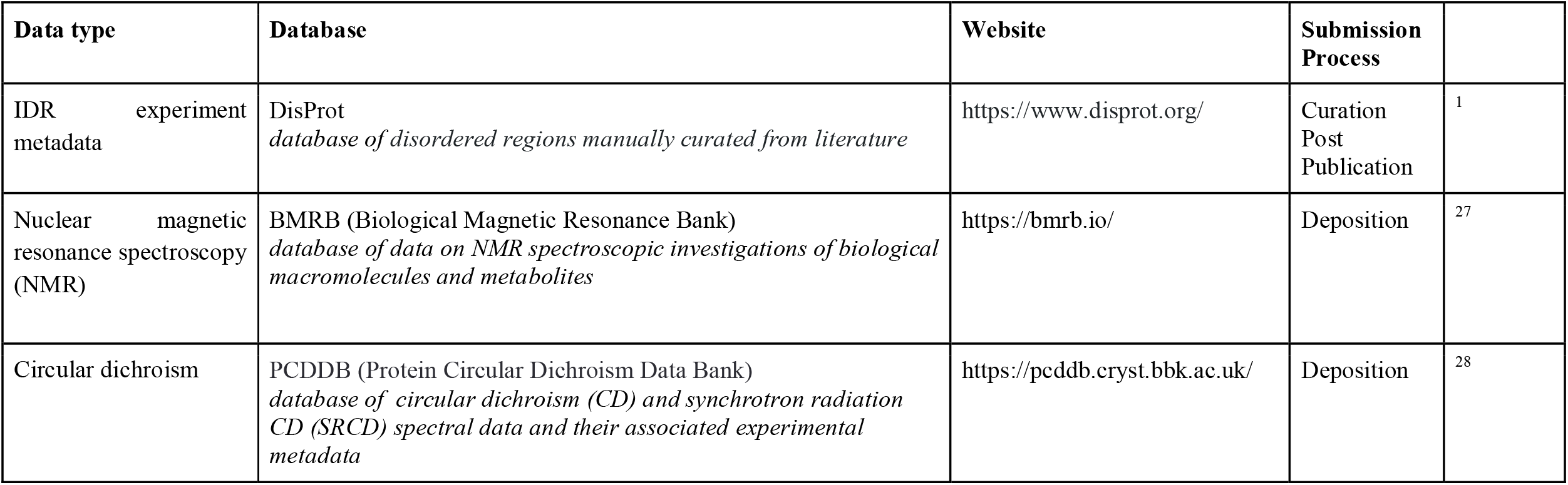
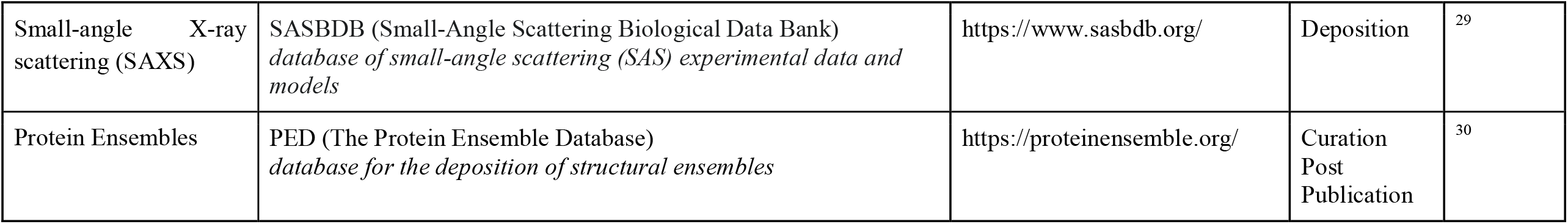
Representative set of databases for the submission of IDR experimental metadata and data.

| Data type | Database | Website | Submission Process |  |
| --- | --- | --- | --- | --- |
| IDR experiment<br>metadata | DisProt<br>*database of disordered regions manually curated from literature* | https://www.disprot.org/ | Curation<br>Post<br>Publication | <sup>1</sup> |
| Nuclear magnetic<br>resonance spectroscopy<br>(NMR) | BMRB (Biological Magnetic Resonance Bank)<br>*database of data on NMR spectroscopic investigations of biological*<br>*macromolecules and metabolites* | https://bmrb.io/ | Deposition | <sup>27</sup> |
| Circular dichroism | PCDDb (Protein Circular Dichroism Data Bank)<br>*database of circular dichroism (CD) and synchrotron radiation*<br>*CD (SRCD) spectral data and their associated experimental*<br>*metadata* | https://pcddb.cryst.bbk.ac.uk/ | Deposition | <sup>28</sup> |
| Small-angle X-ray scattering (SAXS) | SASBDB (Small-Angle Scattering Biological Data Bank)<br>*database of small-angle scattering (SAS) experimental data and models* | https://www.sasbdb.org/ | Deposition | <sup>29</sup> |
| Protein Ensembles | PED (The Protein Ensemble Database)<br>*database for the deposition of structural ensembles* | https://proteinensemble.org/ | Curation<br>Post<br>Publication | <sup>30</sup> |

## Discussion

The IDP community has evolved rapidly in the past 10 years and as methods and technologies have advanced there has been a clear transition in the complexity of the analyses being performed (Figure 1C). This revolution has not been reflected by advances in the data standardisation of the field. Consequently, at all levels there is a requirement to improve the description, curation, storage and dissemination of the fundamental data from these analyses. Guidelines to unambiguously define the key information from an experiment are a crucial first step to simplify data capture, minimise key data loss and standardise data transfer.

Data capture should as much as possible have the flexibility to cover historical experimental approaches, approaches that are cutting edge today, and future advances. The MIADE guidelines store information on the experimental level to allow data to be reinterpreted in the future. While adding experimental parameters and sample components can add considerably to the curation burden they also allow for more nuanced observations to be captured. As IDP experiments become increasingly complex by studying the modulatory effects of proteoforms, concentrations, conditions and binding partners, it is imperative that these rich data on the context of the studied protein region are captured. Furthermore, the attributes that are measured by these approaches are consistently growing, changing from a historically binary order/disorder structure definition to quantitative measures that include dynamics, secondary structure propensity and compaction. This adds a new level of requirements to be considered by comprehensive reporting guidelines. Consequently, the MIADE guidelines will need to evolve over time based on community requirements.

The argument against standardised reporting guidelines has always been the unbalanced burden placed on the reporter. However, the advantages to report far outweigh the effort. Simple accessible data reporting allows relevant data to be easily identified, recovered and reused. Basic administrative advantages include improved data management, minimised data loss associated with group member turnover and simplification of data sharing within and between groups. Refactoring data as method independent metadata allows data to be aggregated but also to be analysed in subsets based on data quality (Figure 1D). Furthermore, data aggregation across complementary methods simplifies cross-validation of data permitting quality to be defined by consensus. Finally, improved data management, in parallel with upgrades to data deposition processes, will improve data transfer to community resources thereby accelerating the open science efforts of the community.

The MIADE guidelines are only an initial step towards standardised and lossless IDP data transfer within the biological community. Three key developments are still required: standardised exchange formats for reporting IDP metadata and raw data, simplified pre- and post-publication data deposition mechanisms for the IDP data repositories, and a community wide agreement to deposit data. The diversity of the methodologies and data in the IDP community has to date proved to be a barrier to data collection. However, standardised guidelines for shared high level metadata annotation such as MIADE will allow the key data to be collected and aggregated across the field. In parallel, each experimental approach in the field can develop method-specific storage and exchange formats and standards for raw data. Enforcing data deposition is a more complex process, however, pressure at the point of publication by journals and reviewers can drive compliance.

It is important that data producers, curators and database developers in the IDP field are conscious of the expanding interest in IDRs by the wider biological community. The growing understanding of the functional significance of IDPs by researchers outside the IDP field has increased the importance of making high quality and understandable IDP data accessible to non-experts such as cell biologists studying the function of IDRs, computational biologists developing tools to analyse IDRs and curators transferring IDR data into community resources.

## Acknowledgements

This work was funded by: ELIXIR, the research infrastructure for life-science data; a Cancer Research UK Senior Cancer Research Fellowship (N.E.D.: C68484/A28159); Carlsberg Foundation Distinguished Fellowship (CF18-0314); Danmarks Grundforskningsfond (DNRF125); National Research, Development and Innovation (NRDI) Fund Young researchers’ excellence programme research grant (project FK128133 to R.P.); European Union’s H-2020 MSCA-RISE programme (grant agreement No. 778247 ‘IDPFun’); EM is supported by Fondazione Umberto Veronesi; the Italian Ministry of University and Research (MUR), PRIN 2017 under grant agreement No 2017483NH8. The authors would like to thank Julie Forman-Kay for her feedback on the MIADE guidelines and the manuscript.

## Notes

### Competing Interest Statement

The authors have declared no competing interest.

